# Phospholipids Uniquely Modify Secondary Structure of α-Synuclein Oligomers

**DOI:** 10.1101/2021.01.15.426884

**Authors:** Tianyi Dou, Lei Zhou, Dmitry Kurouski

## Abstract

Parkinson disease (PD) is a severe neurological disorder that affects more than a million people in the U.S. alone. A hallmark of PD is the formation of intracellular α-synuclein (α-Syn) protein aggregates called Lewy bodies (LBs). Although this protein does not have a particular localization in the central neural system, α-Syn aggregates are primarily found in certain areas of midbrain, hypothalamus and thalamus. Microscopic analysis of LBs revealed fragments of lipid-rich membranes, organelles and vesicles. These and other pieces of experimental evidence suggest α-Syn aggregation can be triggered by lipids. In this study, we used atomic force microscope Infrared (AFM-IR) spectroscopy to investigate structural organization of individual α-Syn oligomers grown in the presence of two different phospholipids vesicles. AFM-IR is a modern optical nanoscopy technique that has single-molecule sensitivity and sub-diffraction spatial resolution. Our results show that α-Syn oligomers grown in the presence of phosphatidylcholine have distinctly different structure than oligomers grown in the presence on phosphatidylserine. We infer that this occurs because of specific charges adopted by lipids, which in turn governs protein aggregation. We also found that protein to phospholipid ratio makes a substantial impact on the structure of α-Syn oligomers. These findings demonstrate that α-Syn is far more complex than expected from the perspective of structural organization of oligomeric species.

## Introduction

α-Synuclein (α-Syn) is the major component of Lewy bodies (LBs), cytoplasmic deposits of proteins in neurons located in several areas of midbrain, hypothalamus and thalamus.^*1-4*^ These α-Syn deposits are a pathogenic hallmark of all synucleinopathies, including Parkinson’s disease (PD), dementia with Lewy bodies (DLB), and multiple system atrophy (MSA).^*1, 5*^ Microscopic analysis of Lewy bodies revealed presence of α-Syn fibrils, long unbranched protein aggregates.^*6, 7*^ *In vitro* experiments confirmed that α-Syn spontaneously aggregate forming first small oligomers that later propagate into fibrils.^*8-12*^ The complexity of cellular environment has strong effect in shaping α-Syn structure: difference in condition may lead to different conformation of α-Syn aggregates.^*13*^ There is a growing body of evidence suggesting that oligomers rather than fibrils are responsible for the onset of PD and the spread of DLB in the brain.^*14 15-18*^ Cryo-electron microscopy (cryo-EM) and solid-state nuclear magnetic resonance (ss-NMR) allowed for elucidation of structural organization of α-Syn fibrils.^*19-23*^ However, the transient nature of oligomers and their morphological heterogeneity limit the use of cryo-EM and ss-NMR for determination of the oligomers’ structure.

AFM-IR is modern analytical technique that can be used to resolve secondary structure of protein oligomers with sub-diffraction spatial resolution.^*24-29*^ Recently reported experimental evidence suggest that AFM-IR has single-monolayer^*30*^ and even single-molecule^*31*^ sensitivity. AFM-IR has been utilized to investigate various topics in biology and surface chemistry including amyloid fibrils,^*25, 26, 32-35*^ plant epicuticular waxes^*36, 37*^, polymers^*38*^, malaria blood cells^*39*^, meteorites^*40*^, bacteria^*41-43*^, liposomes^*44*^ and polycrystalline perovskite films^*45*^. Our group showed that structural characterization of individual α-Syn oligomers can be unravelled using atomic force microscope infrared spectroscopy (AFM-IR).^*15*^ Using AFM-IR, we found that α-Syn aggregation in a lipid-free environment yields structurally diverse oligomers.^*15*^ Some of these oligomers had primarily α-helical/unordered structure, whereas others were mostly composed of antiparallel- and parallel-β-sheet. We also observed structural rearrangement of antiparallel β-sheet to parallel-β-sheet secondary when oligomers propagate into fibrils.^*15*^

Manheim and co-workers recently reported that LBs contain fragments of lipid-rich membranes, organelles and vesicles.^*46, 47*^ Independently, Galvagnion found that rates of α-Syn aggregation can be significantly altered by lipids.^*48-50*^ Moreover, rates of α-Syn aggregation depend on protein-to-lipid-ratio (P:L ratio). It has been shown that an increase in the concentration of dimyristoyl phosphatidylserine (DMPS) relative to concentration of protein first leads to an increase in the rate of α-Syn aggregation. This can be explained from a perspective of lipids as nucleation sites. The larger the number of lipid vesicles the larger the number of α-Syn aggregation sites and thus faster kinetics of protein aggregation. However, a subsequent increase in the amount of lipids causes a decrease in kinetics of protein aggregation. This suggests that delocalization of monomeric α-Syn along surfaces of DMPS vesicles minimizes protein-protein interactions that are necessary for the aggregate formation. These experimental findings suggest that phospholipids may not only alter rates of α-Syn aggregation, but also can change the secondary structure of oligomers. In this study, we used AFM-IR to investigate the structural organization of individual α-Syn oligomers grew in the presence of DMPC (α-Syn-DMPC) at 1:2 (α-Syn-(1:2)-DMPC) and 1:10 (α-Syn-(1:2)-DMPC) ratios. We also utilized AFM-IR to probe the structure of α-Syn oligomers grown in the presence of DMPS (α-Syn-DMPS) at 1:2 (α-Syn-(1:2)-DMPS) and 1:10 (α-Syn-(1:2)-DMPS) ratios due to high abundancy of both of these phospholipids in cell and organelles membranes or neurons.

## Result and Discussion

We found that oligomers at the early stage of α-Syn aggregation (day 2) had spherical morphology with heights ranging from 1 to 10 nm, Figure 1. Specifically, most of α-Syn-(1:2)-DMPC and α-Syn-(1:10)-DMPC oligomers had 1-3 nm and 1-6 in height, respectively. α-Syn-(1:2)-DMPS and α-Syn-(1:10)-DMPS were 1-4 and 1-5 nm respectively. These results suggest that all α-Syn-DMPS and α-Syn-DMPC oligomers exhibit high similarity in both topology and size.

**Figure 1.**
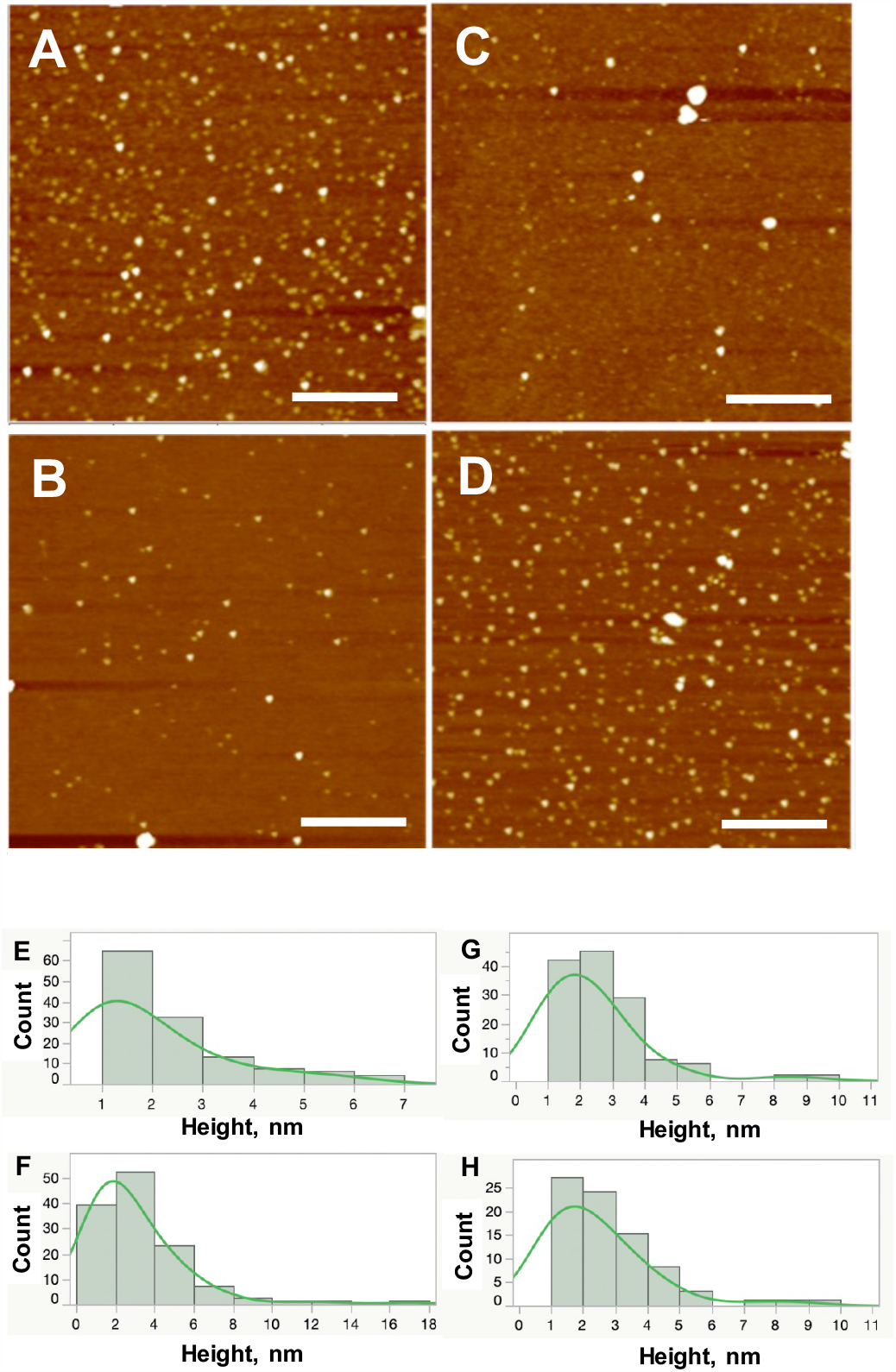
AFM images (A-D) and corresponding height profiles (E-H) of α-Syn-(1:2)-DMPC (A, E); α-Syn-(1:10)-DMPC (B, F); α-Syn-(1:2)-DMPS (C, G); α-Syn-(1:10)-DMPS (D, H). Scale bars are 1 micron.

AFM-IR analysis of individual α-Syn-(1:2)-DMPC, α-Syn-(1:10)-DMPC, α-Syn-(1:2)-DMPS and α-Syn-(1:10)-DMPS oligomers revealed major differences in their structural organization. Specifically, α-Syn-(1:2)-DMPC aggregates exhibited intense vibrational bands at 1624-1632 cm^-1^ which could be assigned to the parallel β-sheet, Table 1, Figure 2. At the same time, vibrational bands at 1672-1682 and 1702 cm^-1^, which could be assigned to β-turns and anti-parallel β-sheet respectively, had weak intensities. These results show that both parallel- and antiparallel-β-sheet, as well as β-turns are present in α-Syn-(1:2)-DMPC oligomers.

**Table 1.** Vibrational bands and their assignments for phospholipids and proteins.

| Band wavenumber ($\text{cm}^{-1}$) | Vibration mode | Assignment |
| --- | --- | --- |
| 866-950 | $\nu(\text{C-N}^+-\text{C})$ | lipid<sup>51</sup> |
| 974 | $\nu(\text{C-N}^+-\text{C})$ | lipid<sup>51</sup> |
| 1020-1028 | C-O stretch | lipid<sup>52, 53</sup> |
| 1050-1100 | $\nu(\text{PO}_2^-)$ | lipid<sup>51</sup> |
| 1160-1200 | P=O; C-O-P | lipid<sup>54</sup> |
| 1256-1264 | $\nu(\text{PO}_2^-)$ | lipid<sup>55</sup> |
| 1289-1298 | N-H |  |
| 1350 | Stretching C-O | lipid<sup>56</sup> |
| 1380 | $\text{CH}_3$, C-O stretching | lipid<sup>51, 55, 56</sup> |
| 1610-1640 | amide I | β-sheet<sup>15</sup> |
| 1640-1653 | amide I | random coil<sup>15</sup> |
| 1648-1660 | amide I | α-Helix<sup>15</sup> |
| 1660-1685 | amide I | $3_{10}$-Helix, type II β-turn<sup>15</sup> |
| 1675-1697 | amide I | Antiparallel β-sheet<sup>15</sup> |
| 1730-1766 | $\nu(\text{C=O})$ | lipid<sup>56</sup> |

**Figure 2.**
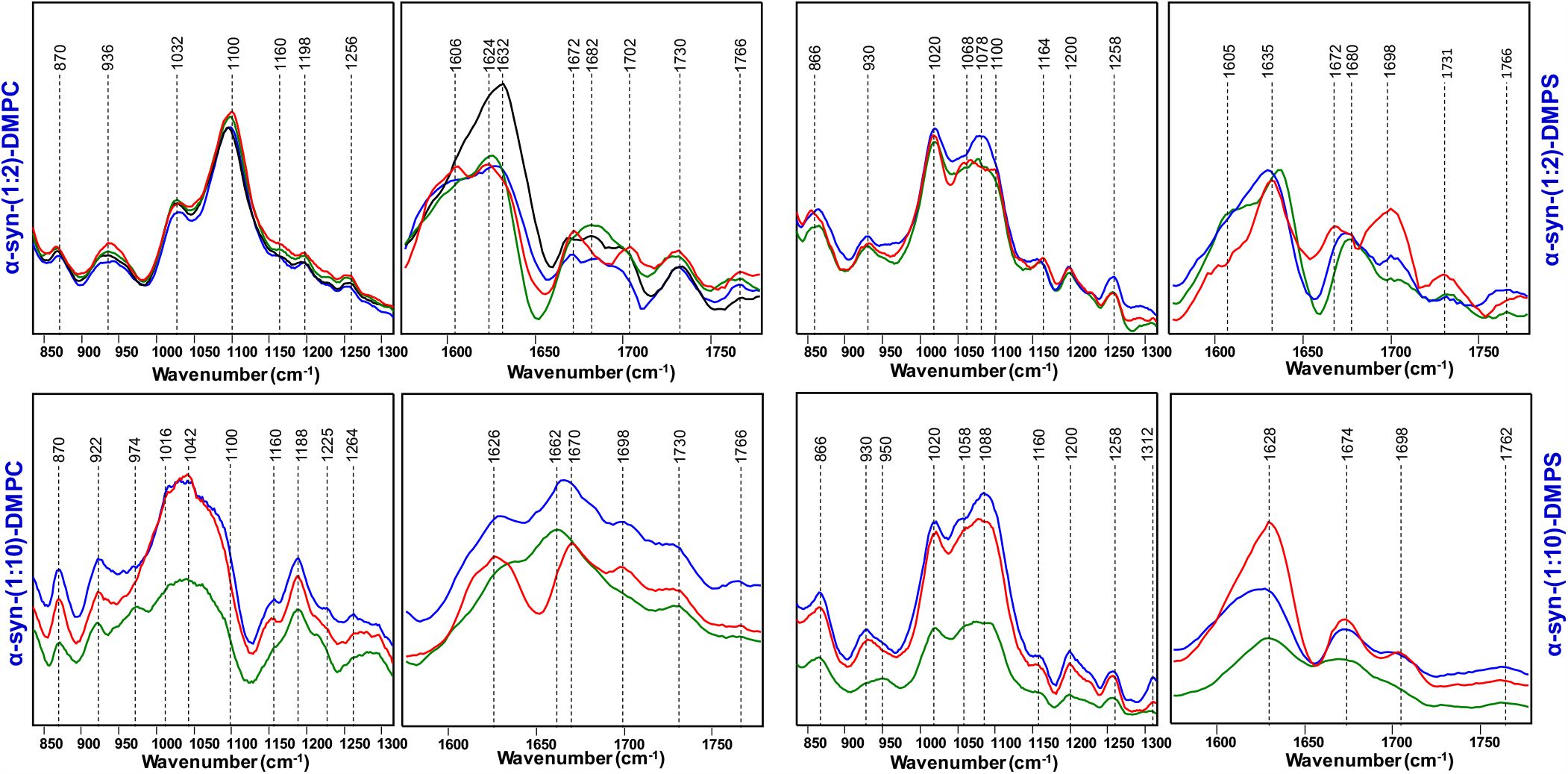
AFM-IR spectra of individual α-Syn-(1:2)-DMPC, α-Syn-(1:10)-DMPC, α-Syn-(1:2)-DMPS and α-Syn-(1:10)-DMPS oligomers. Blue, red, green correspond to 1,2,3 and black corresponds to 4 in Figure 3.

Our experimental findings suggest that structural organization of α-Syn-(1:10)-DMPC and α-Syn-(1:2)-DMPC oligomers is substantially different. AFM-IR spectra collected from α-Syn-(1:10)-DMPC oligomers exhibited vibrational bands centered at 1662 and 1670 cm^-1^ indicating high content of α-helix/unordered protein secondary structure in these oligomers. We also found vibrational bands at 1626 and 1698 cm^-1^ that demonstrate the presence of both parallel and anti-parallel β-sheet in the structure of α-Syn-(1:10)-DMPC oligomers. However, the relative content of these secondary structures is drastically different between α-Syn-(1:2)-DMPC and α-Syn-(1:10)-DMPC oligomers. In α-Syn-(1:2)-DMPC oligomers, the content of parallel β-sheet was nearly three times higher than the amount of unordered and antiparallel β-sheet, whereas in α-Syn-(1:10)-DMPC oligomers the amount of α-helix/unordered protein secondary structure was nearly twice higher than the amount of parallel and antiparallel β-sheet.

AFM-IR spectra of both α-Syn-(1:2)-DMPC and α-Syn-(1:10)-DMPC oligomers exhibited vibrational bands that could be assigned to DMPC, Figure 2. Specifically, we found vibrational band at 870, 922-936 cm^-1^, which could be assigned to C-N^+^-C vibrations of this phospholipid. We also observed vibrations at 1016-1032 cm^-1^, which could be assigned to C-O stretch of the phospholipid and vibrational bands at 1042-1100 cm^-1^ that originate from the phosphor group vibrations. Lastly, in the spectra of both α-Syn-(1:2)-DMPC and α-Syn-(1:10)-DMPC, we found a band at 1256-1264 cm^-1^, which originates form PO2^-^ vibration, as well as vibrational bands at 1730 and 1766 cm^-1^, which could be assigned to carbonyl vibration of DMPC. These findings indicate that phospholipid is present in the structure of both α-Syn-(1:2)-DMPC and α-Syn-(1:10)-DMPC. In addition to the change in the intensity of vibrational bands, we found a shift of PO_2_^-^ vibration in the spectra of α-Syn-(1:2)-DMPC and α-Syn-(1:10)-DMPC. In the spectrum of α-Syn-(1:2)-DMPC, this vibration is centered at 1100 cm^-1^, whereas in the spectrum of α-Syn-(1:10)-DMPC this band was found at 1042 cm^-1^. This blue shift suggests that in the structure of α-Syn-(1:2)-DMPC, the charged phosphor group is located in the hydrophilic environment, whereas in α-Syn-(1:10)-DMPC the phosphor-group is squeezed to the aqueous environment. Although extensive calculations are required to fully elucidate conformations of DMPC in both α-Syn-(1:2)-DMPC and α-Syn-(1:10)-DMPC oligomers, one can conclude that this phospholipid has different organization in those two classes of oligomers.

We also found that α-Syn-(1:10)-DMPC oligomers exhibit very similar if not identical structural organization. This conclusion can be made based on similar profiles of amide I vibrations (1622-1704 cm^-1^). The higher intensity of the spectrum of one of the oligomers (blue trace) comparing to the other three (black, red and green traces) is likely due to differences in the height of these oligomers. The higher oligomer exhibited stronger AFM-IR spectrum comparing to smaller oligomers. Although α-Syn-(1:2)-DMPC oligomers exhibit similar spectra, their structural heterogeneity is substantially larger. Specifically, we found that one of examined oligomers (black trace) exhibited much stronger intensity of 1622-1632 cm^-1^ bands comparing to the other three (green, blue and red). This evidence suggests that this oligomer had higher amount of parallel β-sheet comparing to the other oligomers. Also, small changes in intensities of vibrational bands at 1672-1683 cm^-1^ (β-turns) and 1696-1704 cm^-1^ (antiparallel-β-sheet) suggest about small deviations in the amount of these secondary structural elements in their structure. Thus, α-Syn-(1:2)-DMPC are not as structurally homogeneous as α-Syn-(1:10)-DMPC oligomers.

Spectroscopic analysis of α-Syn-DMPS oligomers revealed the presence of vibrational bands that can be assigned to parallel (1620-1635 cm^-1^) and antiparallel β-sheet (1694-1696 cm^-1^), Figure 2. These spectra also exhibited vibrations at 1672-1680 cm^-1^ that could be assigned to β-turn secondary structure. Based on these results, we can conclude that both α-Syn-(1:2)-DMPS and α-Syn-(1:10)-DMPS are primarily composed of parallel, antiparallel β-sheet and β-turns. We also found that the α-Syn-(1:2)-DMPS exhibit more heterogeneous content of antiparallel-β-sheet comparing to α-Syn-(1:10)-DMPS oligomers. Specifically, the heterogeneity arises from different ratios of parallel and antiparallel β-sheet in analysed oligomers. In two of the analysed α-Syn-(1:2)-DMPS oligomers, parallel β-sheet prevailed over the content of antiparallel β-sheet. However, in the third oligomer, the amount of parallel and antiparallel β-sheet was nearly identical.

AFM-IR showed that α-Syn-DMPS oligomers exhibit vibration bands that could be assigned to DMPS. Similar to DMPC, vibrational bands at 866, 930 cm^-1^ could be assigned to C-N^+^-C bond in the lipid. The vibration at 1020 cm^-1^ could be assigned to C-O stretch, whereas bands from 1058 to 1100 cm^-1^ could be assigned to the phosphate group vibration. Both α-Syn-(1:2)-DMPS and α-Syn-(1:10)-DMPS exhibited high similarity of the lipid region of the spectrum. This suggests about similar interaction patterns between α-Syn and DMPS under two different P:L ratios.

We observed some heterogeneity within both α-Syn-(1:2)-DMPS and α-Syn-(1:10)-DMPS. α-Syn-(1:2)-DMPS oligomers exhibited heterogeneity related to different ratios of parallel *vs* antiparallel-β-sheet (red vs green and blue curves, α-Syn-(1:2)-DMPS, Figure 2). The same conclusions can be made for α-Syn-(1:10)-DMPS. We found that ratio of parallel vs antiparallel-β-sheet (black *vs* red and blue curves) change form one oligomer to another. Nevertheless, spectroscopic analysis of both α-Syn-DMPS and α-Syn-DMPC oligomers revealed substantially less structural heterogeneity comparing to α-Syn aggregates grown in the lipid-free environment.^*15*^ This findings point out on the efficient nucleation of α-Syn aggregation by both DMPS and DMPC.^*57*^ Also, based on the discussed above results, we can conclude that early state α-Syn oligomers grown in the presence of DMPC and DMPS have drastically different structures comparing to α-Syn oligomers grown in the lipid-free environment.

Next, we performed imaging of these oligomers using AFM-IR. Images were collected at 1100 cm^-1^ (phospholipids), 1624 cm^-1^ (parallel-β-sheet), 1655 cm^-1^ (α-helix/unordered protein secondary structure) and 1694 cm^-1^ (antiparallel-β-sheet) to probe localization lipids and protein secondary structure elements in the structure of all four types of oligomers (Figure 3). We found that both phospholipids are homogeneously distributed in α-Syn-(1:2)-DMPC, α-Syn-(1:10)-DMPC, α-Syn-(1:2)-DMPS and α-Syn-(1:10)-DMPS. AFM-IR imaging confirmed discussed above ratio of parallel-to-antiparallel β-sheet in α-Syn-(1:2)-DMPC and α-Syn-(1:10)-DMPC. Specifically, α-Syn-(1:10)-DMPC exhibit very similar ratios of parallel and antiparallel β-sheets in their structure (Figure 3, F and H), whereas parallel β-sheet dominates in the structure of α-Syn-(1:2)-DMPC. We also found that α-Syn-(1:2)-DMPC are nearly entirely composed of parallel β-sheet (Figure 3, C), whereas a substantial amount of α-helix has been observed in α-Syn-(1:10)-DMPC (Figure 3, G). α-Syn-(1:2)-DMPS and α-Syn-(1:10)-DMPS exhibited primarily parallel β-sheet secondary structure, with detectable α-helix content at the edges of the oligomers.

**Figure 3.**
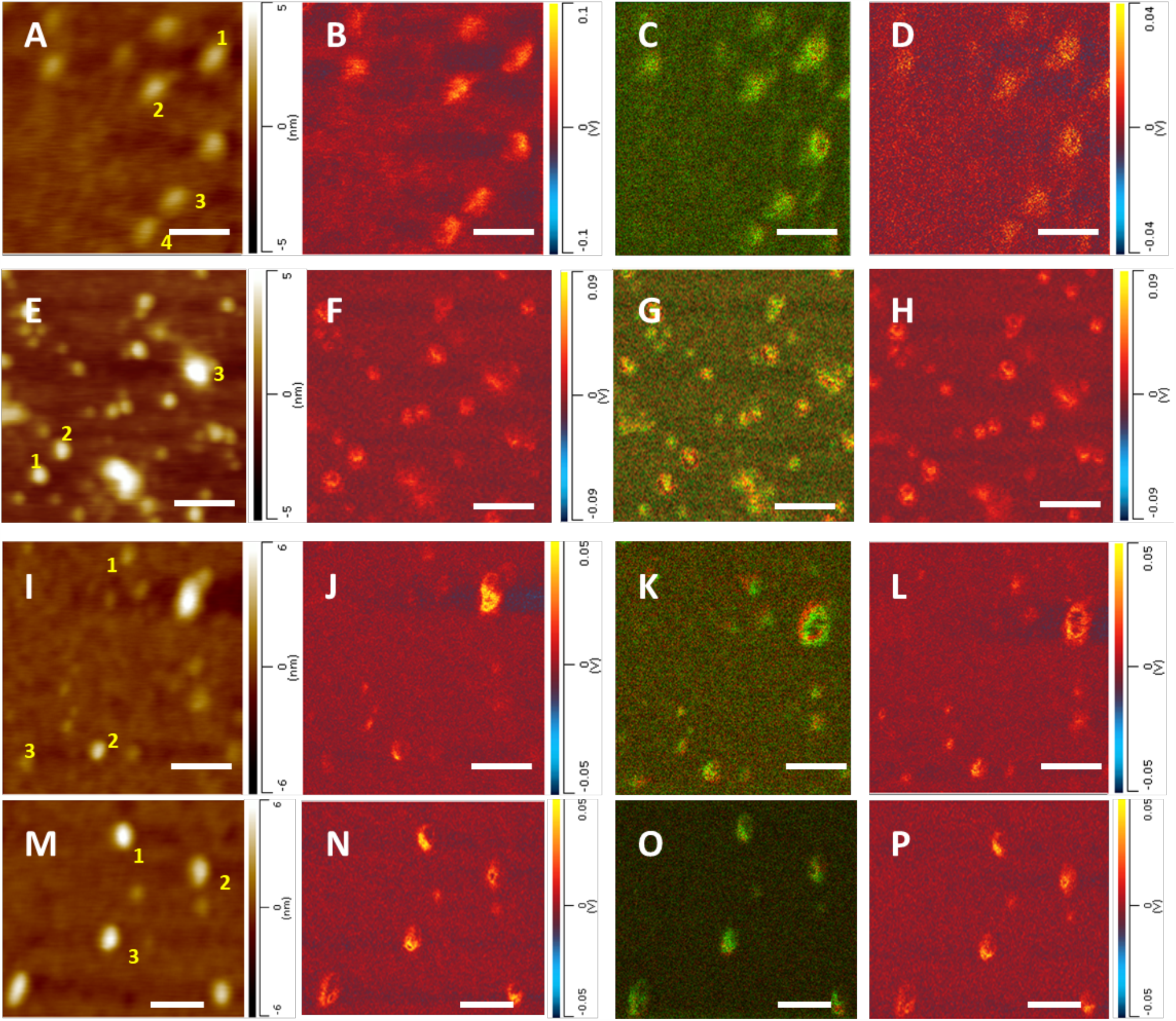
AFM-IR chemical map of α-Syn-(1:2)-DMPC (A-D), α-Syn-(1:10)-DMPC (E-H), α-Syn-(1:2)-DMPS (I-L) and α-Syn-(1:10)-DMPS oligomers (M-P). AFM height images (A, E, I, M), IR absorption map at 1100 cm^-1^ (B, F, J, N), IR ratio overlaying maps of 1624cm^-1^ (green) / 1655cm^-1^ (red) (C, G, K, O) and IR absorption map at 1694 cm^-1^ (D, H, L, P). Scale bar, 100nm.

Numerous evidence indicates the toxicity of the oligomers, which could result from the interaction with lipids and disruption of membranes.^*14*^ Danzer et.al used different incubation conditions identified three types of oligomers. Interestingly, these different types of oligomers have distinct cellular effects, though all toxic.^*58*^ Chen et.al also found oligomers with antiparallel β-sheet rich structure possessed stronger toxicity compared with parallel rich fibrils^*29*^. These findings indicate that the distinct properties of different oligomers might because of the structure difference. Using AFM-IR, the heterogeneous oligomer structure can all be identified. Our findings might help to better understand the underlying cause of oligomer toxicity.

## Conclusions

Our findings demonstrate that α-Syn oligomers grown in the presence of DMPC and DMPS have significantly different secondary structures comparing to α-Syn oligomers grown in the lipid-free environment. Moreover, both phospholipids present in the structure of the oligomers. We also found that α-Syn-DMPC oligomers are drastically different from α-Syn-DMPS aggregates in terms of the possessed secondary structure. Lastly, we found that not only the chemical structure of the lipid, but also P:L ratio alters the secondary structure of α-Syn oligomers.

## Methods

### Protein and lipids preparation

The preparation of lipid vesicles in this work was similar to that reported previously by Galvagnion et al^*59*^. The lipids 0.6 mg of DMPS and DMPC (Avanti Polar Lipids Inc.) were separately dissolved in 2.6ml of HEPES buffered saline (HBS) pH 7.4. The lipid vesicle, LUVs were prepared using sonication (Fisher Scientific, Ultrasonic Bath, 3 × 5 min). Sonication was performed in ice water bath. After centrifugation, the sizes of lipid LUVs were checked using dynamic light scattering (DLS). The size of DMPC LUVs was shown to consist of a distribution centering at 60nm and 100nm, while the size of DMPS LUVs consists of the distribution centering at 110 nm.

The preparation of α-Synuclein aggregates is followed from our previous work from Zhou and Kurouski.^*15*^ α-Syn (AnaSpec, CA, USA) was dissolved to the final concentration of 50 μM, in 50nM Tris buffer, 150 mM NaCl, pH at 7.5. Next, the solution of the α-Syn was immediately mixed with either DMPC or DMPS lipid vesicles solution and kept at room temperature without agitation. The final concentration of α -Synuclein is 160 μM.

### AFM-IR

The α-Synuclein aggregates were deposited on silicon wafer. AFM-IR imaging was conducted using a Nano-IR3 system (Bruker, Santa Barbara, CA, USA). The IR source was a QCL laser. Contact-mode AFM tips (ContGB-G AFM probe, NanoAndMore) were used to obtain all spectra and maps. IR maps at 1624, 1655, 1694 cm^-1^ wavenumber values were obtained to study the secondary structure of α-Syn-DMPS/DMPC oligomers, 1100 cm^-1^ to study the presence of these phospholipids in α-Syn aggregates (Table 1). AFM height and deflection images were acquired simultaneously with IR maps. Spatial resolution of AFM-IR was ∼ 10nm. Totally, 20 point spectra were taken from every analysed oligomer. The spectra were filtered by Deglitch and Savitzky-Golay in Analysis Studio software. The parameter of Savitzky-Golay is 2 polynomial order and 7 side points.

## Associated Content

### Supporting Information Available

Supporting information includes atomic force microscopy height image and dynamic light scattering results of DMPS and DMPC lipid vesicles; Averaged AFM-IR spectra of α-synuclein-1:2-DMPC, α-synuclein-1:10-DMPC, DMPC Luvs, α-synuclein-1:2-DMPS, α-synuclein-1:10-DMPS and DMPS Luvs.

## Acknowledge and Acknowledgements

The work was supported by the start-up funds from Texas A&M University.

## Conflicts of interest

The authors declare no competing financial interest.

---

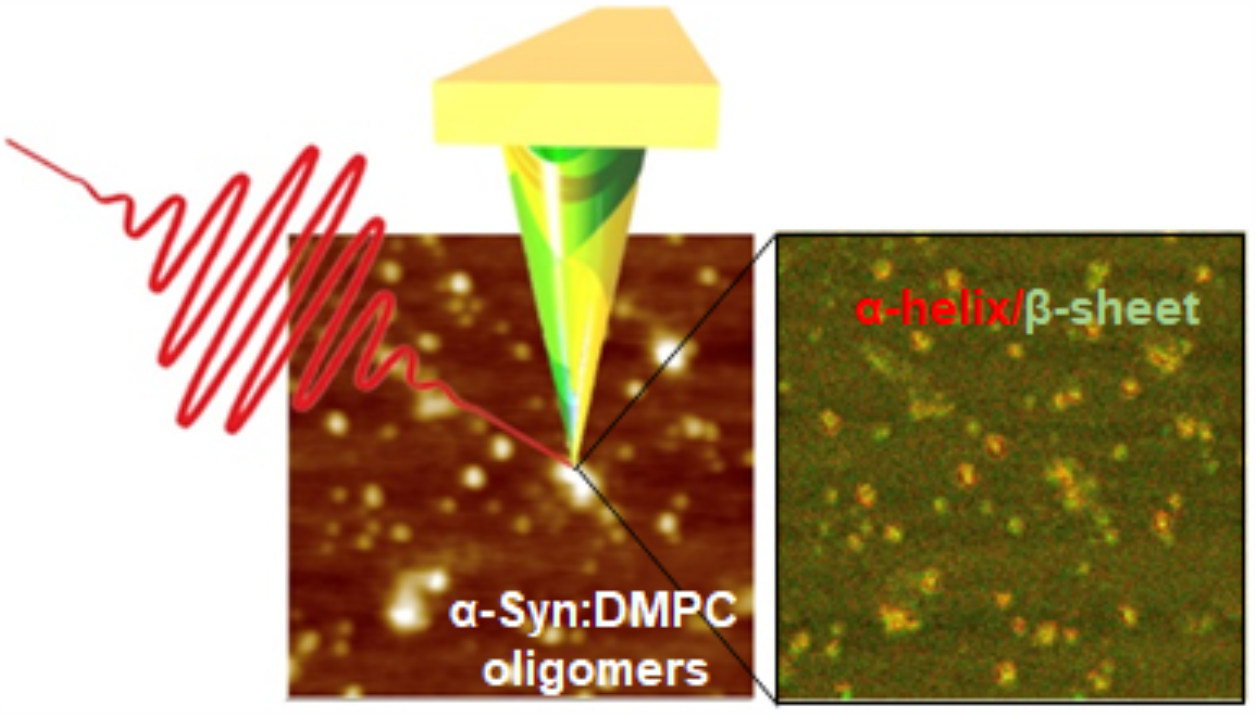

## Notes

### Competing Interest Statement

The authors have declared no competing interest.

