## Supplemental files for "Phospholipids Uniquely Modify Secondary Structure of α-Synuclein Oligomers"

**Contents:**

**Figures:**

**Figure S1: AFM height image and DLS of DMPS lipid vesicle**

**Figure S2: AFM height image and DLS of DMPC lipid vesicle**

**Figure S3: Averaged AFM-IR spectra of α-synuclein-1:2-DMPC, α-synuclein-1:10-DMPC and DMPC Luvs**

**Figure S4: Averaged AFM-IR spectra of α-synuclein-1:2-DMPS, α-synuclein-1:10-DMPS and DMPS Luvs**

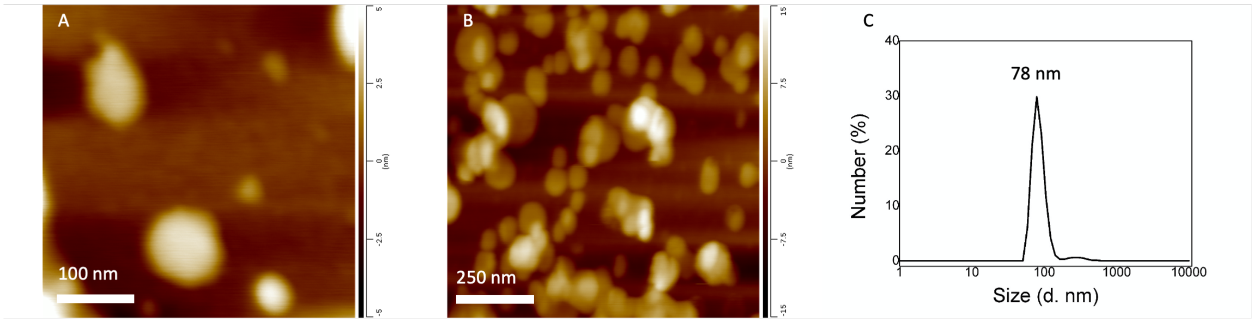


Figure S1. AFM height image of DMPS lipid vesicle (A and B). Dynamic light scattering (DLS) reveals the size of DMPS lipid vesicle (C).

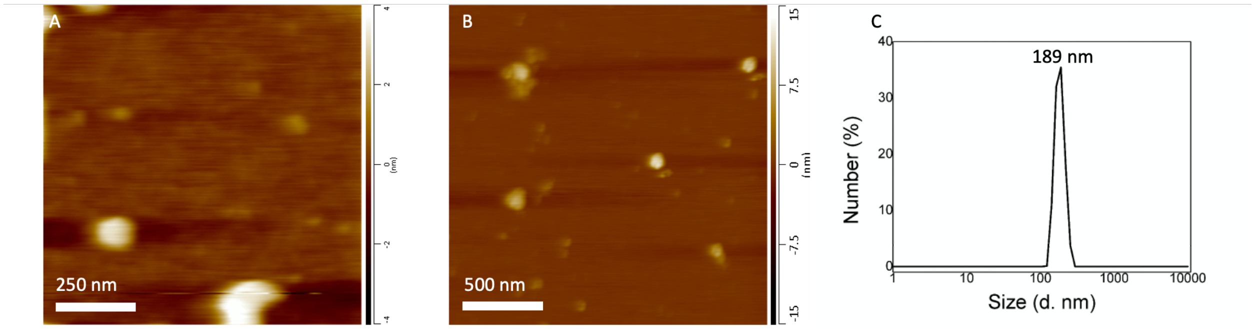


Figure S2. AFM height image of DMPC lipid vesicle (A and B). Dynamic light scattering (DLS) reveals the size of DMPC lipid vesicle (C).

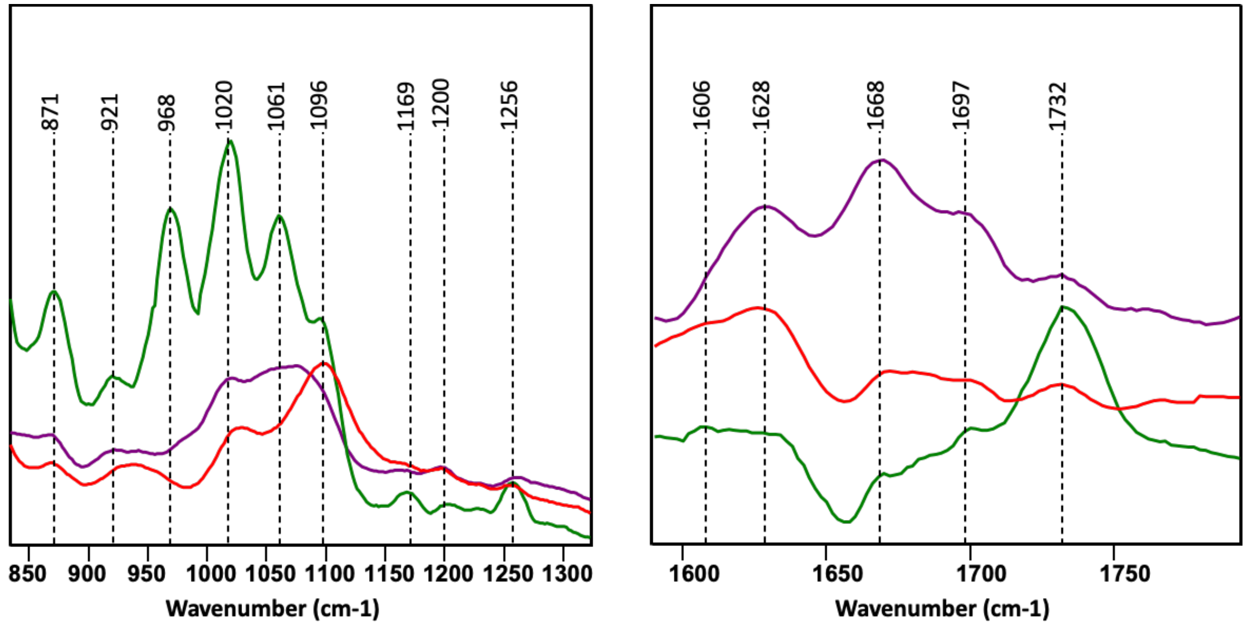


Figure S3. Averaged AFM-IR spectra of α-synuclein-1:2-DMPC (red), α-synuclein-1:10-DMPC (purple) and DMPC Luvs(green) on lipid (800-1350 $cm^{-1}$) and amide I region (1550-1800 $cm^{-1}$).

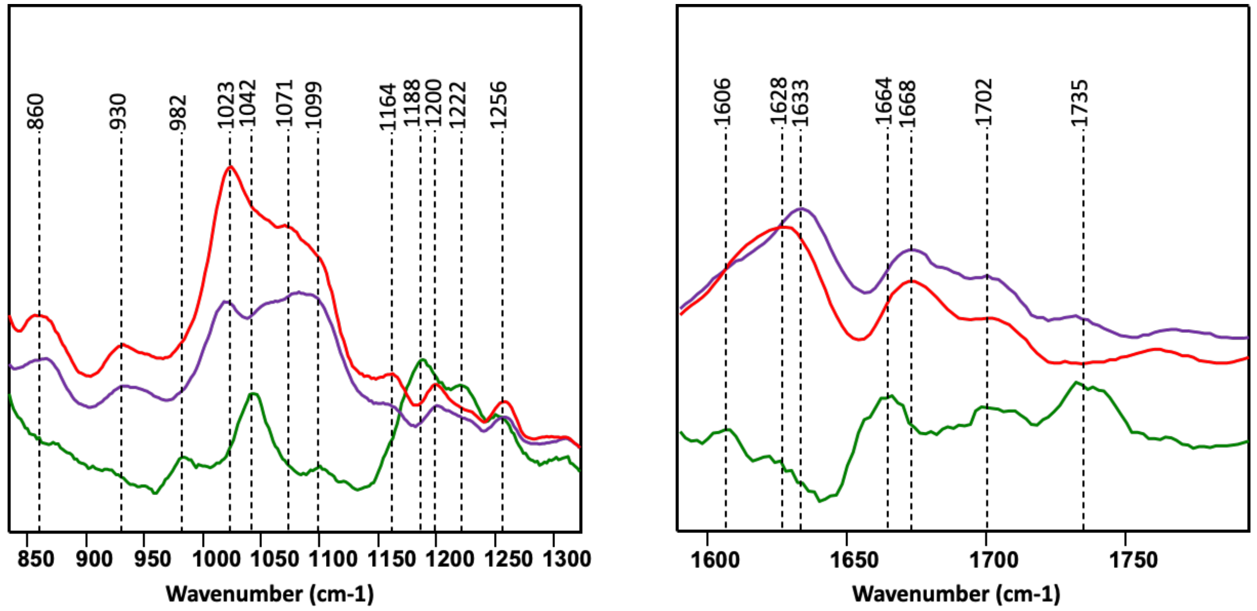


Figure S4. Averaged AFM-IR spectra of α-synuclein-1:2-DMPS (red), α-synuclein-1:10-DMPS (purple) and DMPS Luvs(green) on lipid (800-1350 $cm^{-1}$) and amide I region (1550-1800 $cm^{-1}$).
